# Prolonged stalling of RNA Polymerase II at DNA damage explains phenotypical differences between Cockayne and UV-sensitive syndromes

**DOI:** 10.1101/2023.05.17.541110

**Authors:** Camila Gonzalo Hansen, Barbara Steurer, Roel C. Janssens, Di Zhou, Marjolein van Sluis, Hannes Lans, Jurgen A. Marteijn

## Abstract

Faithful transcription of eukaryotic genes by RNA polymerase II (Pol II) is essential for proper cell function. Nevertheless, the integrity of the DNA template of Pol II is continuously challenged by different sources of DNA damage, such as UV-light, that impede transcription. When unresolved, these transcription-blocking lesions (TBLs) can cause cellular dysfunction, senescence and apoptosis, eventually resulting in DNA damage-induced aging. Cells counteract these deleterious effects by Transcription-Coupled Nucleotide Excision Repair (TC-NER), which specifically removes TBLs, thereby safeguarding transcription. TC-NER initiation relies on the concerted actions of the CSB, CSA and UVSSA proteins, and loss of either of these factors results in a complete TC-NER deficiency. Although their TC-NER defect is similar, UVSSA loss results in UV-Sensitive Syndrome (UV^S^S), with only mild phenotypes like freckling and photosensitivity, while loss of CSA or CSB activity results in the severe Cockayne Syndrome (CS), characterized by premature aging, progressive neurodegeneration and mental retardation. Thus far the underlying mechanism for these striking differences in phenotypes remains unclear. Using live-cell imaging approaches, here we show that in TC-NER proficient cells lesion-stalled Pol II is swiftly resolved by repair of the TBL. However, in CSA and CSB knockout (KO) cells, elongating Pol II remains chromatin-bound. This lesion-stalled Pol II will obstruct other DNA transacting processes and will also shield the damage from repair by alternative pathways. In contrast, in UVSSA KO cells, Pol II is removed from the TBL by VCP-mediated proteasomal degradation, thereby, allowing alternative repair mechanisms to remove the TBL.

## Introduction

Unperturbed transcription of eukaryotic genes by RNA polymerase II (Pol II) is crucial for proper cell function. However, the DNA strand transcribed by Pol II is continuously threatened by agents of both endogenous and exogenous origins that cause damage that can severely impede or even completely block transcription (1, 2). Endogenous transcription-blocking DNA damage can result from a variety of sources such as oxidative stress and aldehydes (3, 4), but also transcription-replication conflicts or DNA-secondary structures may impede transcription (1, 5). Well-known examples of exogenous DNA damaging agents that can induce transcription-blocking lesions (TBLs) are chemical mutagens and ultra-violet radiation (UV) (3), of which the latter induces helix-distorting DNA lesions that block the forward progression of Pol II on DNA (6). Unresolved TBLs can lead to a transcription impediment or transcription of aberrant RNA (7), causing cellular dysfunction, senescence and apoptosis, eventually resulting in damage-induced aging (2, 8). To overcome these severe consequences, TBLs are efficiently removed by Transcription-Coupled Nucleotide Excision Repair (TC-NER), thereby safeguarding transcription (2, 9, 10).

Lesion-stalled Pol II is recognized by CSB (11), which uses its ATP-dependent translocase activity to discriminate between lesion-stalled Pol II and naturally-paused Pol II (12, 13). The prolonged CSB-binding to Pol II is hypothesized to result in the recruitment of CSA (14). CSA is part of the Cullin 4-based CRL4^CSA^ E3-ligase, which upon activation by NEDD8 conjugation, ubiquitylates both CSB (15) and Pol II (16). Ubiquitylation of specifically the K1268 residue of RPB1, the largest subunit of Pol II (16, 17), which is stimulated by ELOF1, drives efficient recruitment of the downstream TC-NER factors UVSSA and TFIIH (16, 18–20). Due to its interaction with the de-ubiquitylating enzyme USP7, UVSSA inhibits CSB ubiquitylation thereby preventing its degradation (21). In addition, UVSSA recruits TFIIH (14, 22). After TFIIH binding to the TBL, XPA is recruited, which stimulates the DNA damage verification activity of TFIIH (23) and the recruitment and orientation of the ERCC1/XPF and XPG nucleases that finally excise the TBL (24).

The biological relevance of TC-NER is best illustrated by the early onset of Cockayne syndrome (CS) in patients with mainly inactivating mutations in the CSA and CSB genes (25). CS patients suffer from growth and development complications appearing within their first year of life, microcephaly and progressive neurological dysfunction, photosensitivity, and symptoms of accelerated aging such as hearing loss, ophthalmological degeneration, osteoporosis, liver and kidney failure and motor abnormalities. Due to the severity of the disease, the average life expectancy of CS patients is only 12 years (26, 27). Strikingly, the phenotypes observed in UV-sensitive syndrome (UV^S^S) patients, which have an equal TC-NER deficiency at the cellular level as CS patients, are remarkably milder (28, 29). Except for specific mutations in for example CSA (30), the main gene mutated in UV^S^S patients is UVSSA (21, 31, 32). UV^S^S patients have mild cutaneous phenotypes including photo-hypersensitivity and freckling without the neurodegeneration and other severe phenotypes of CS (33). Thus far the underlying mechanism for this striking difference in phenotypes remains unclear, which may be due to the fact that most research is performed in rare non-isogenic patient cells.

The observed phenotypical differences have been linked to additional functions of CS proteins compared to UVSSA. Several functions for CS proteins, mainly for CSB, outside of TC-NER were described, including preservation of mitochondrial function (34, 35), regulation of transcription mainly for neuronal development (36) and control of redox balance (37). However, the exact roles of CSA and UVSSA in these processes remain much less understood. Instead, the progressive nature of the CS phenotype is in line with the stochastic accumulation of endogenous DNA damage or toxic repair intermediates in TC-NER deficient cells, suggesting that a defective genome maintenance mechanism is the common denominator of this TC-NER linked disorder (2). An interesting observation is that deficiencies in the first initiation steps of TC-NER are characteristic of CS, due to mutations in CSA and CSB, while defective downstream steps are characteristic of UV^S^S, due to UVSSA mutations. Mostly, in CS cells, the activity of the CRL4^CSA^ E3-ligase is impeded, for example due to reduced CSA binding caused by mutations at the CSB-CSA interface, or by mutations severely compromising CSA structure (13, 25, 38, 39). As lesion-stalled Pol II is one of the key substrates of CRL4^CSA^ (16), it will not be resolved by TC-NER in CS cells. We therefore hypothesize that lesion-stalled Pol II will remain chromatin-bound for a prolonged time as its degradation is hampered due to the impeded CRL4^CSA^ activity. This chromatin-bound Pol II will not only create a toxic persistent transcription block, which will most likely also impede other DNA transacting processes, but will also shield the TBL that is buried within Pol II from alternative repair pathways. On the contrary, in UV^S^S cells in which the CRL4^CSA^ E3 ligase is still active (30), lesion-stalled Pol II can still be degraded without prior repair of the lesion. This CRL4^CSA^-mediated degradation of lesion-stalled Pol II will eventually allow the access of alternative repair pathways to resolve the TBL. We postulate that this differential processing of Pol II could explain the difference in phenotypes between CS and UV^S^S.

To study the differential chromatin binding of Pol II in TC-NER deficient cells we generated isogenic knock-ins (KIs) of endogenously GFP-tagged Pol II in knockout (KO) cells of the different TC-NER factors (40). This allowed us to study Pol II chromatin binding upon DNA damage using live-cell imaging approaches and cell fractionation assays (41). We show that in TC-NER proficient cells lesion-stalled Pol II is swiftly resolved by repair of the TBL. In line with our hypothesis, we find that while in TC-NER deficient CSA and CSB KO cells lesion-stalled elongating Pol II remains chromatin-bound, while in UVSSA KO cells Pol II is removed from the damage by VCP-mediated proteasomal degradation. Together, our data shows that the prolonged stalling of Pol II at TBLs correlates with the CS phenotype, and suggests that the failure to remove toxic TC-NER intermediates will cause persistent transcription stress. Additionally, these intermediates will shield TBLs from alternative repair pathways. Therefore, prolonged stalling of Pol II at a TBL will contribute to cellular functional decline and increased apoptosis and senescence, which are the basis of the severe, progeroid phenotype of CS.

## Results

### Differential Pol II chromatin-binding upon DNA damage in WT, CS and UV^S^S cells

To test whether the phenotypic differences in CS and UV^S^S could be explained by differential chromatin-binding dynamics of Pol II upon TBL induction we employed Fluorescence Recovery after Photo-bleaching (FRAP) (42) of GFP-tagged Pol II, using knock-in (KI) cells expressing a GFP-tagged version of RPB1 at endogenous levels. FRAP of GFP-RPB1 KI cells is a sensitive tool to study the Pol II dynamics in living cells, at the different stages of the transcription cycle in unperturbed conditions (40) and upon DNA damage (19, 41). We generated isogenic GFP-RPB1 KIs (MRC5^GFP-RPB1^) in CSB, CSA, UVSSA, XPA and XPC knock-out (KO) cell lines. CSB and CSA KO cells represent CS and UVSSA KO cells represent UV^S^S. XPC KO cells were included as a control for defective Global Genome NER (GG-NER), while XPA KO cells are deficient in both GG-NER and TC-NER. We use this cell line as a control, because Pol II is expected to be already evicted from TBLs when XPA is normally recruited during TC-NER. KO of NER proteins was confirmed by western blotting (Fig. 1A). UVSSA KO was confirmed by genotyping (Fig. S1) as endogenous UVSSA could not be detected by western blot. Full TC-NER deficiency in these KOs was tested by clonogenic cell survival (Fig. 1B) and by measuring their ability to restart transcription after UV damage (Fig. 1C-D). WT cells showed proficient colony formation and transcription restart. A similar UV-sensitivity and loss of transcription restart was observed in CSB, CSA and UVSSA KO cell lines. XPA KO cells showed a slight increase in UV-sensitivity as these cells are deficient for both GG-NER and TC-NER. XPC KO cells were as expected sensitive to UV in the colony formation assay due to their GG-NER deficiency, but showed transcription restart. Importantly, these data demonstrate a similar TC-NER deficiency in CSB, CSA and UVSSA KO cells.

**Figure 1.**
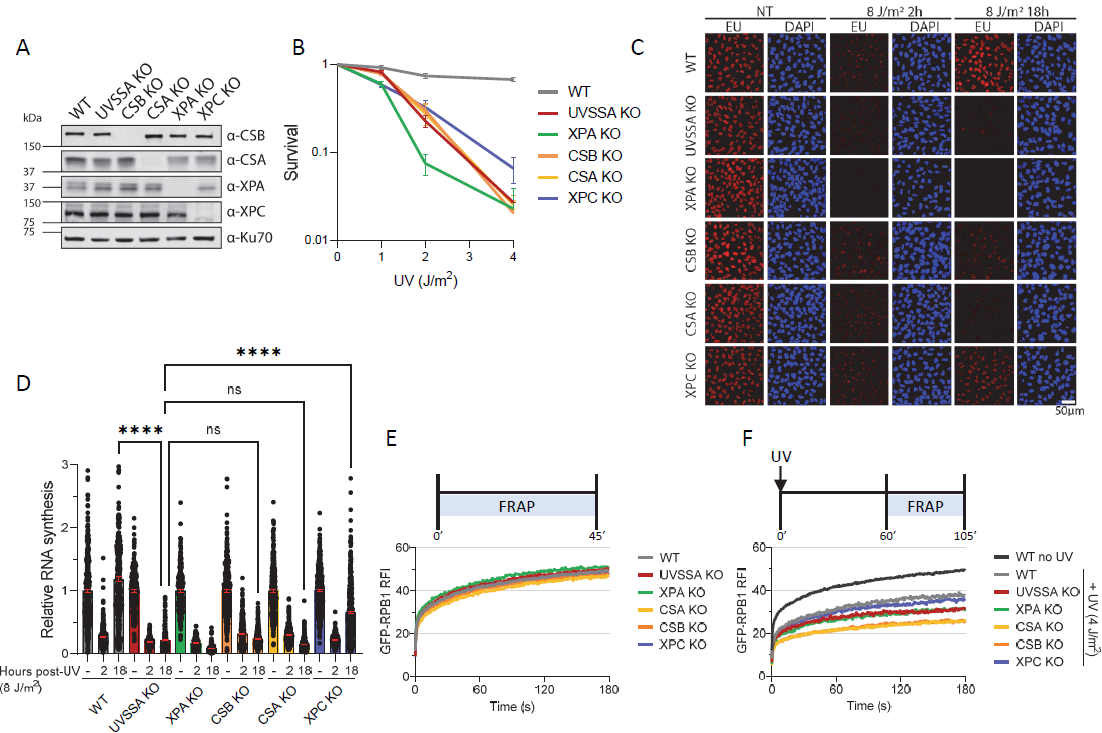
Different Pol II chromatin-binding dynamics after DNA damage observed in WT, CS and UV^S^S cells. A. Western Blot analysis using the indicated antibodies of MRC5^GFP-RPB1^ WT cells and CRISPR/Cas9-mediated knock out (KO) cells of the indicated TC-NER proteins CSA and CS B, of the GG-NER protein XPC, and of the XPA protein involved in both sub-pathways. Ku70 was used as a loading control. Endogenously expressed UVSSA could not be detected and UVSSA KO was assessed by genotyping (Fig. S1A). B. Relative clonogenic survival of MRC5^GFP-RPB1^ WT or specified KO cells exposed to the indicated doses of UV (J/m^2^). ± SEM, n = 2. C. Representative images of transcription levels as determined by relative EU incorporation in irradiated and non-irradiated cells 2h and 18h after UV exposure (8 J/m^2^). NT = Non-treated. D. Quantification of transcription restart after UV damage as determined by relative EU incorporation in MRC5^GFP-RPB1^ WT and indicated KO cells at the specified time points after UV exposure (8 J/m^2^) as shown in (**C**). The relative integrated fluorescence intensity was normalized to mock-treated levels and set at 1. Columns indicate average relative integrated fluorescence intensity and error bars ± SEM. n ≥ 178 cells per condition from at least 2 independent experiments. ns = not significant, ****P< 0.0001. E. Fluorescence Recovery after photo-bleaching (FRAP) analysis of GFP-RPB1 in unperturbed conditions in WT or the indicated KO MRC5^GFP-RPB1^ cells. GFP-RPB1 was bleached and fluorescence intensity was measured every 0.4 sec for 3 min, background-corrected and normalized to pre-bleach fluorescence intensity, which was set to 100. Average Relative Fluorescence Intensity (RFI) of n ≥ 16 cells per condition from at least 2 independent experiments. F. FRAP analysis similar as in (**E**), but 1-2h after UV (4 J/m^2^) irradiation. GFP-RPB1 mobility of unperturbed WT cells is plotted in black for comparison. Average RFI of n ≥ 21 cells per condition from at least 3 independent experiments.

Next, we used FRAP of GFP-RPB1 to compare Pol II chromatin binding kinetics in unperturbed conditions or after inducing TBLs by UV irradiation. As promoter-bound Pol II complexes are chromatin-bound on average for less than a minute, they mostly contribute to the fast, initial recovery of fluorescence (<50 sec) during GFP-RPB1 FRAP. In contrast, elongating Pol II complexes are chromatin-bound for >20 min on average, and are therefore mainly represented by the later part of the GFP-RPB1 FRAP curve (>100 sec) (40). GFP-RPB1 FRAP analysis showed highly similar Pol II chromatin-binding kinetics in WT and the different KO cells in unperturbed conditions (Fig. 1E), indicating that transcription is not affected in unperturbed conditions in these KO cells. Irradiation with a low dose of 4 J/m^2^ UV significantly reduced Pol II mobility in WT cells, evident from the reduced fluorescence recovery compared to unperturbed cells (Fig. 1F). This immobilization affected both the initial part of the FRAP curve (<50 sec) as well as the later part, evident from the reduced slope of the FRAP curve at time points >100 sec. In line with previous findings (41), this can be explained by (i) degradation of promoter-bound Pol II which happens in a TC-NER independent manner and by (ii) stalling of elongating Pol II at TBLs. As expected, TC-NER proficient XPC KO cells, showed Pol II chromatin-binding dynamics comparable to WT cells. In TC-NER deficient CSB, CSA, UVSSA and XPA KO cells, however, Pol II was further immobilized. This was especially evident by the reduced slope of the FRAP curve at the later part of the curve, mainly representing elongating Pol II, suggesting that Pol II elongation rates were reduced, most likely caused by accumulation of lesion-stalled Pol II. Interestingly, this UV-induced Pol II immobilization was increased in CSA and CSB KO cells compared to UVSSA and XPA KO cells (Fig. 1F). These data suggest that either elongating Pol II is bound for a longer time, or more elongating Pol II is chromatin-bound, in the absence of CSB or CSA. This implies that CSA and CSB contribute to the release of lesion-stalled Pol II in the absence of TC-NER.

### Elongating Pol II is longer bound to TBLs in the absence of CSA and CSB

Elongating Pol II molecules that are trailing, or halted, behind a lesion-stalled Pol II, may conceal the difference in residence times of lesion-stalled Pol II complexes in the different TC-NER KO cells. This is especially important since the number of lesion-stalled Pol II complexes will be rather small compared to the total number of elongating Pol II molecules at the relatively low, but physiological relevant damage loads used. To circumvent this, we developed an approach to more specifically study the residence time of lesion-stalled Pol II. To do so, after DNA damage infliction we blocked *de novo* transcription initiation using the CDK7 inhibitor THZ1 (43) to reduce the number of elongating Pol II molecules trailing behind lesion-stalled Pol II (Fig. 2A). We found that 45 min transcription inhibition resulted in an almost complete mobilization of Pol II in undamaged conditions (Fig. 2B). The increased Pol II mobilization can be explained by the accumulation of freely diffusing, *i.e*. highly mobile, Pol II, caused by a loss of immobile, chromatin-bound elongating Pol II. This immobile Pol II is lost due to transcription termination at the end of genes in combination with a loss of new transcription initiation events (40). The Pol II mobilization upon THZ1 treatment in unperturbed conditions happened to similar extents in WT and TC-NER KO cells. To study the residence time of lesion-stalled Pol II, we first UV irradiated the cells and allowed 30 min for elongating Pol II to encounter a lesion, thereafter *de* novo transcription initiation was inhibited by THZ1. THZ1 treatment in UV-irradiated WT cells resulted in a full mobilization of elongating Pol II similar to unperturbed conditions (Fig. 2C). This indicates that under these conditions lesion-stalled Pol II is efficiently resolved, most likely due to TC-NER. In contrast, Pol II immobilization was increased in UV-irradiated CSB and CSA KO cells compared to WT cells (Fig. 2C-D), showing that 2 h after DNA damage induction ∼10-20% of all Pol II molecules in these cells remained stalled at the TBL. While a prolonged stalling of only 10-20% of all Pol II complexes may seem like a rather mild consequence, with ∼50.000 Pol II molecules present in the cell (40), these represent ∼5.000-10.000 large macromolecular chromatin-bound complexes that will interfere with most DNA transacting processes in the gene bodies, an important part of the genome. Interestingly, THZ1 treatment following UV-induced DNA damage in XPA and UVSSA KO cells led to Pol II TBL-binding kinetics highly resembling those of WT cells (Fig. 2C-D). This indicates that within this 75-120 min time-period lesion-stalled Pol II is removed from the chromatin, even in the absence of TC-NER. Furthermore, this finding suggests that the intermediate Pol II immobilization observed without THZ1 upon UV-irradiation in XPA and UVSSA KO cells (Fig. 1F) is mainly caused by elongating Pol II molecules that are trailing behind the initially stalled Pol II, which will subsequently stall on the same lesion due to the absence of TC-NER.

**Figure 2.**
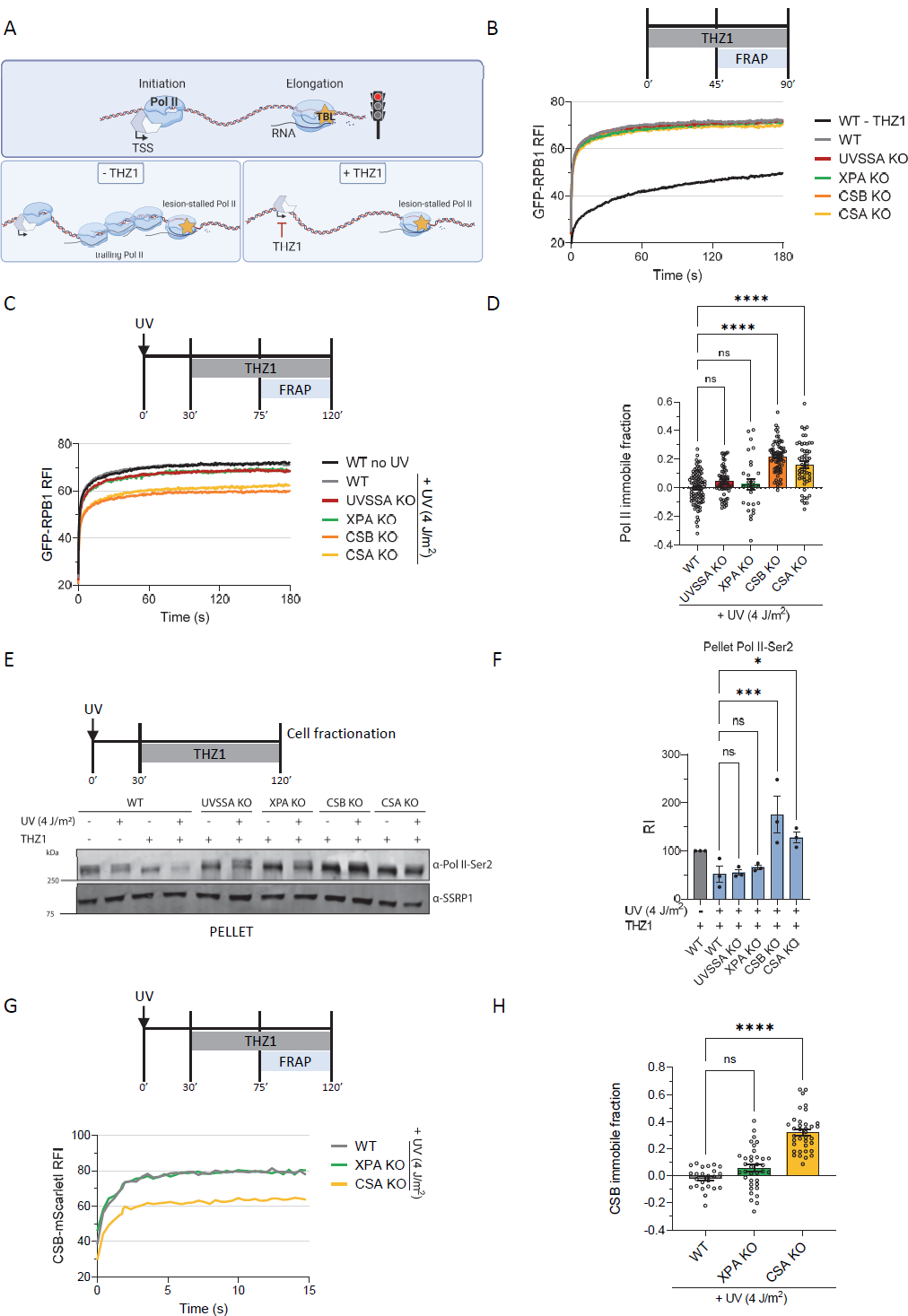
Prolonged chromatin-binding of elongating Pol II upon DNA damage in CSB and CSA KO cells. A. Cartoon of experimental setup to study the residence time of mainly lesion-stalled Pol II at TBLs. Elongating Pol II was allowed to run into UV-induced lesions for 30 min after which the *de novo* transcription initiation was blocked by the CDK7 inhibitor THZ1. This reduced the stalling of elongating Pol II molecules behind lesion-stalled Pol II in a “traffic jam”. This allowed us to mainly study lesion-stalled Pol II chromatin binding, as 45 min after CDK7 inhibition most elongating Pol II will be released from the chromatin due to transcription terminations in unperturbed conditions. TSS = Transcription start site. B. Chromatin binding of Pol II as determined by FRAP of GFP-RPB1 in the indicated MRC5^GFP-RPB1^ WT and TC-NER-deficient cell lines in unperturbed conditions, or upon THZ1 treatment. Average Relative Fluorescence Intensity (RFI) as calculated by normalizing post-bleach values to pre-bleach values which were set at 100. RFI of n ≥ 29 cells per condition from at least 3 independent experiments. C. FRAP analysis of GFP-RPB1 similar to (**B**), but after UV (4 J/m^2^) irradiation 30 min prior to THZ1 treatment. GFP-RPB1 mobility of unperturbed WT cells is plotted in black for comparison. Average RFI of n ≥ 27 cells per condition from at least 3 independent experiments. D. Quantification of GFP-RPB1 immobile fraction after DNA damage obtained from (**B, C**). Immobile fraction in WT and TC-NER KO cells was calculated by comparing RFI after UV to mock-treated sample and normalizing to their own non-irradiated RFI values (See methods for more details). n ≥ 27 cells per condition from at least 3 independent experiments ± SEM. ns = not significant, ****P<0.0001. E. Representative western blot of pellet fraction after cell fractionation in MRC5^GFP-RPB1^ WT and indicated TC-NER KO cells. CTD-Ser2-phosphorylated RPB1 (Pol II-Ser2) staining in chromatin-bound fraction in unperturbed and UV-irradiated (4 J/m^2^) conditions upon THZ1 treatment to quantify chromatin-bound elongating Pol II. SSRP1 is used as loading control. F. Quantification of Pol II-Ser2 signal from 3 independent experiments as in (**E**). Pol II-Ser2 signal was normalized to SSRP1 (loading control) signal and to the mock-treated sample for each cell line and set at 100. RI = Relative intensity, ns = not significant, *P<0.05, ***P<0.001, ± SEM, n = 3. G. FRAP of CSB-mScarletI in HCT116^CSB-mScarletI^ WT and TC-NER KO cells upon UV irradiation (4 J/m^2^) followed by THZ1 treatment. CSB-mScarletI was bleached in a strip across the nucleus and fluorescence intensity was measured every 0.4 sec for 16 sec 0-30 min after UV irradiation with the indicated UV doses. n ≥ 24 cells per condition from at least 2 independent experiments. H. Quantification of CSB-mScarletI immobile fraction after DNA damage obtained from (**G, S2D**). Immobile fraction was calculated as in (**D**). n ≥ 24 cells per condition from at least 2 independent experiments ± SEM. ns = not significant, ****P<0.0001.

To confirm that elongating Pol II is chromatin-bound for a prolonged time upon DNA damage induction in CSA and CSB KO cells, we performed cell fractionation experiments followed by western blot analysis of RPB1. Chromatin-bound elongating Pol II was quantified by staining of serine 2-phosphorylated RPB1 (Pol II-Ser2) in the chromatin fraction (41). As expected, THZ1 treatment resulted in a reduction of elongating Pol II, in the absence of DNA damage. Levels of elongating Pol II were slightly further reduced upon TBL-induction (Fig. 2E-F), which could be explained by Pol II release from the chromatin or by its dephosphorylation. In line with our FRAP results, a similar reduction in chromatin-bound Pol II-Ser2 comparable to WT was found in UVSSA and XPA KO cells upon DNA damage induction. In contrast, Pol II-Ser2 showed increased chromatin-binding in CSB and CSA KO cells upon TBL induction, confirming the prolonged stalling of Pol II in these cells.

In cells, lesion-stalled Pol II is specifically recognized by CSB, resulting in a stable CSB-Pol II interaction (11–13). Therefore, CSB chromatin-binding, as determined by FRAP in CSB-mScarletI KI cells, can be used as a sensitive measure for lesion-stalled Pol II (19). Already at a low UV dose of 2 J/m^2^ a clear CSB immobilization can be observed, which increases in a UV-dose dependent manner (Fig. S2A). To study CSB chromatin-binding dynamics in TC-NER deficient cells, we generated isogenic KOs of CSA (CS) and XPA in HCT116^CSB-mScarletI^ cells (Fig. S2B). TC-NER deficiency was confirmed by clonogenic cell survival (Fig. S2C). UVSSA KO cells could not be used in this assay, as UVSSA is known to be crucial to stabilize CSB upon DNA damage by the recruitment of USP7 (21, 30). No differences in CSB mobility upon THZ1 treatment in the absence of DNA damage were detected between the WT and TC-NER KO cells (Fig. S2D). Next, we tested CSB chromatin-binding upon UV irradiation followed by THZ1 treatment and observed no immobilization in WT cells, which can be explained by TC-NER mediated TBL removal. Similar results were obtained in XPA KO cells, suggesting that lesion-stalled Pol II is removed from the chromatin. However, in CSA KO cells CSB showed a strong immobilization, which corroborates that in the absence of CSA, Pol II is stalled for a prolonged time on chromatin upon DNA damage (Fig. 2G-H). Taken together, these live cell imaging and cell fractionation assays show that Pol II is efficiently removed from TBLs in the absence of functional TC-NER in UVSSA and XPA KO cells. However, this does not happen in CSA and CSB KO cells, resulting in the prolonged chromatin-binding of lesion-stalled Pol II.

### Increased DNA damage-induced Pol II degradation in UVSSA-deficient cells

Next, we set out to study the underlying mechanism explaining how lesion-stalled Pol II is displaced from the chromatin in the absence of TC-NER in UVSSA and XPA KO cells. In UVSSA and XPA KO cells the CRL4^CSA^ E3 ligase is still recruited to lesion-stalled Pol II (14). As CRL4^CSA^ has been shown to stimulate the ubiquitylation of elongating Pol II upon DNA damage (16, 31), Pol II may be degraded in UVSSA and XPA KO cells while in CSA and CSB KO cells this cannot happen due to the absence of CRL4^CSA^. To test this hypothesis, we assessed Pol II degradation by quantifying endogenously expressed GFP-RPB1 protein levels by flow cytometry, which is a sensitive approach to quantify the total cellular Pol II levels without any antibody or extraction bias (40). In conditions in which new transcription initiation events are inhibited by THZ1, exactly as used in the FRAP experiments (Fig. 2), UV-irradiation induced a minor Pol II degradation in WT cells, which was exacerbated in UVSSA KO, but completely absent in CSB and CSA KO cells (Fig. 3A). Importantly, although the difference in Pol II levels in UVSSA KO compared to CSA and CSB KO cells seems rather small (∼10%), this is in line with the ∼10-20% immobilized Pol II observed in CSA and CSB KO cells (Fig. 2D), suggesting that the TBL-stalled Pol II is removed from chromatin by degradation in UVSSA KO cells. Interestingly, while in XPA KO cells Pol II is also released from the chromatin similarly as in UVSSA KO cells, no increased Pol II degradation compared to WT cells could be observed. This indicates that in XPA KO cells Pol II is not degraded but is likely released in a reaction step downstream of UVSSA. XPA is recruited to the TBL after TFIIH binds to the damaged DNA, and Pol II is therefore most likely already removed from the TBL preceding this reaction step, either by backtracking or chromatin release (2). The difference in Pol II degradation in UVSSA and XPA KO cells, suggests that in the absence of TC-NER, Pol II can be released from damaged chromatin by two different mechanisms.

**Figure 3.**
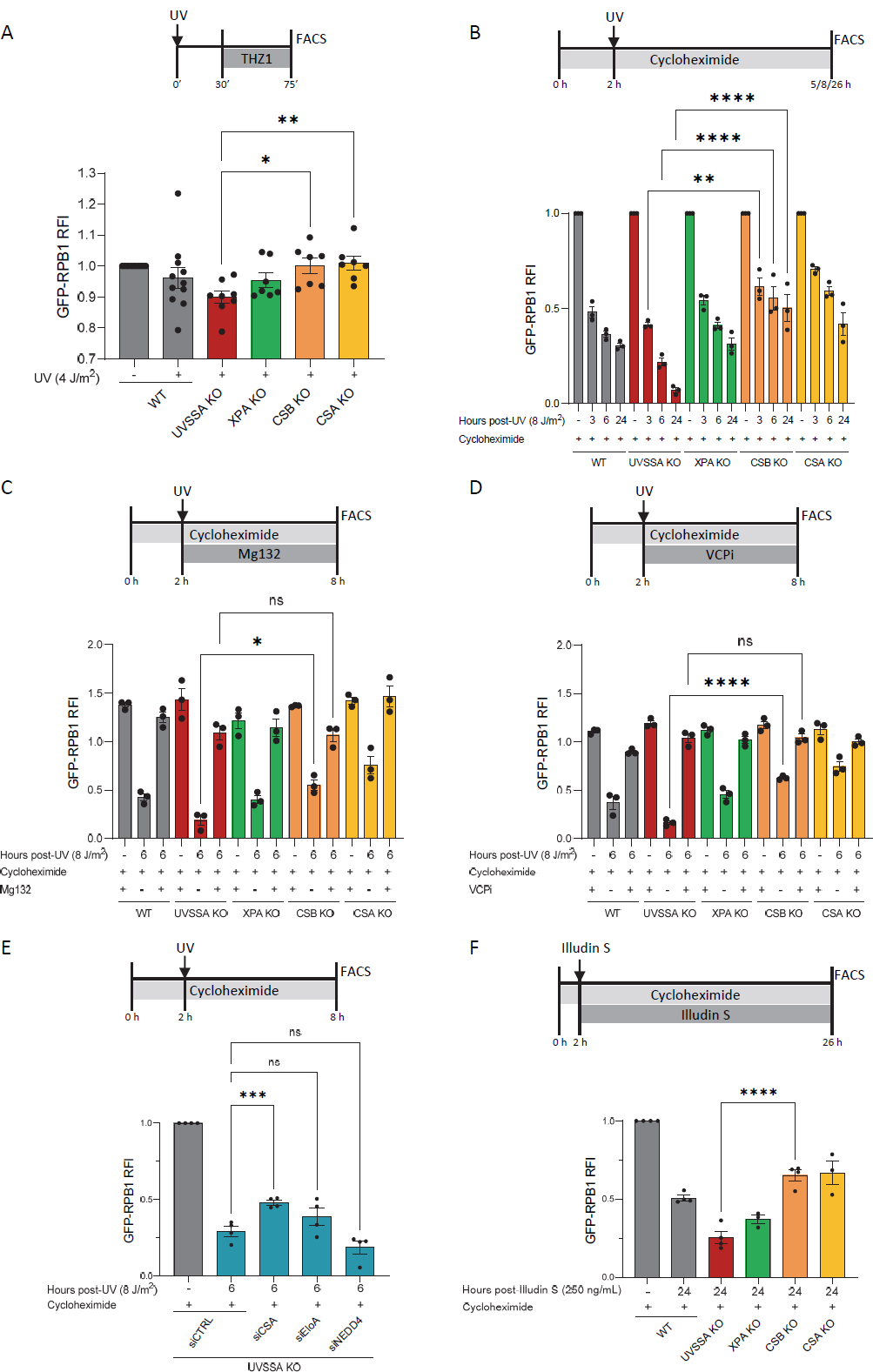
Absence of lesion-stalled Pol II degradation in CSA and CSB KO cells. A. Pol II levels in MRC5^GFP-RPB1^ WT and indicated TC-NER cells were determined by measuring GFP-RPB1 intensity by flow cytometry, measuring at least 10.000 single cells per independent experiment. Cells were irradiated with UV (4 J/m^2^) 30 min prior to THZ1 treatment. GFP-RPB1 levels in UV-irradiated cells were normalized to non-damaged samples for each cell line that were set at 1. Columns represent average RFI of n ≥ 7 independent experiments ± SEM. B. Pol II levels in the indicated MRC5^GFP-RPB1^ cells as in (**A)** at the indicated times after UV-irradiation (8 J/m^2^) in the presence of cycloheximide (100 µM). Cycloheximide was added 2h prior to UV-irradiation and maintained for the duration of the experiment. UV-treated samples were corrected to non-damaged GFP-RPB1 levels of each cell line at each time point and normalized to sample treated for 5 h with cycloheximide which was set at 1. Columns represent average RFI of n = 3 independent experiments ± SEM. C. Pol II levels as in (**B**) with the addition of proteasome inhibition with Mg132 (50 µM) which was added at the time of UV-irradiation (8 J/m^2^). GFP-RPB1 levels in all samples were corrected and normalized to non-damaged samples treated 8 h with cycloheximide without proteasome inhibitor which were set at 1. Columns represent average RFI of n = 3 independent experiments ± SEM. D. Pol II levels as in (**C**) but now in the presence of VCP inhibition using NMS-873 (5 µM). n = 3 independent experiments ± SEM. E. Pol II levels in UVSSA KO cells as in (**B**) upon transfection of the indicated siRNAs and harvested 6 h post-irradiation (8 J/m^2^). siCTRL is a non-targeting siRNA used as control. UV-treated samples were normalized to non-damaged samples treated 8 h with cycloheximide which were set at 1. Columns represent average RFI of n = 4 independent experiments ± SEM. F. Pol II levels upon treatment with Illudin S (250 ng/mL) for 24 h and cycloheximide as in (**B**). Illudin S treated samples were normalized to non-damaged samples treated 26 h with cycloheximide which were set at 1. Columns represent average RFI of n ≥ 3 independent experiments ± SEM.

When studying Pol II degradation in the presence of THZ1, we mostly assessed the stability of the first Pol II that was stalled at a TBL, as successive rounds of Pol II stalling at unrepaired TBLs were prevented due to inhibition of transcription initiation. However, when lesion-stalled Pol II is degraded, consecutive rounds of Pol II stalling at unrepaired TBLs might result in successive rounds of Pol II degradation. Therefore, we quantified Pol II degradation over a period of 24 h after UV in the absence of THZ1. In these conditions a ∼50% loss of Pol II was observed in WT cells (Fig. S3A). However, these measurements were obscured by increased Pol II levels 24 h after UV-irradiation, most likely due to *de novo* Pol II synthesis upon transcription restart after TBLs are repaired. To study specifically Pol II degradation without considering effects of *de novo* translation, protein synthesis was inhibited with the ribosome inhibitor cycloheximide. In these conditions a gradual drop in Pol II levels upon UV damage was observed in WT cells (Fig. 3B). Of note, in these conditions where *de novo* translation is present, Pol II degradation is observed in all cell lines, especially at the earliest measured time-point (3h post-damage). This indicates that in addition to the CSA and CSB dependent degradation of lesion-stalled Pol II (Fig. 3A), also a TC-NER independent degradation of Pol II is observed, which could for example be attributed to the recently described degradation of promoter-bound Pol II, which happens shortly after UV-induced DNA damage independently of TC-NER (41). In addition to this effect, a gradual Pol II degradation could be observed in XPA KO cells similar as in WT cells, confirming that in XPA KO cells Pol II is degraded to a similar level. In CSA and CSB KO cells Pol II degradation was much less pronounced, which is consistent with the impeded degradation of lesion-stalled Pol II. In sharp contrast, we found an strong increased Pol II degradation over time in UVSSA KO cells, with almost no Pol II left after 24 h, which is likely caused by consecutive rounds of Pol II encountering unrepaired TBLs followed by its degradation.

As Pol II degradation correlates with the presence of the CRL4^CSA^ E3 ligase, this suggests that Pol II is being degraded in a ubiquitin and 26S proteasome dependent manner. To test this, we assessed Pol II levels upon proteasome inhibition using Mg132. Proteasome inhibition rescued DNA damage-induced Pol II loss in the different TC-NER KO cells to a similar extent (Fig. 3C), indicating that Pol II is being degraded by the 26S proteasome. Importantly, this also indicates that the additional loss of Pol II in UVSSA KO cells compared to CSA and CSB KO cells is caused by proteasomal degradation of lesion-stalled Pol II. As the extraction of chromatin-bound Pol II upon DNA damage has been described to depend on the ubiquitin selective segregase p97/VCP (41, 44), we tested its involvement in this process. Similarly as upon proteasome inhibition, VCP inhibition by NMS-873 led to a comparable rescue of Pol II levels in all cell lines (Fig. 3D), implying that in UVSSA KO cells lesion-stalled Pol II is ubiquitylated to be subsequently extracted from the chromatin and degraded. Therefore, we explored the contribution of the CRL4^CSA^ E3 ligase to the Pol II degradation observed in UVSSA KO cells, and compared this to the contribution of the Elongin A (EloA) and NEDD4 E3 ligase complexes, that have been described to ubiquitylate elongating Pol II as part of the last-resort Pol II degradation pathway (45). While depletion of NEDD4 or EloA did not significantly reduce Pol II degradation in UVSSA KO cells, depletion of CSA clearly reduced Pol II degradation compared to control conditions, suggesting that indeed CRL4^CSA^ is the main contributor to Pol II ubiquitylation in UVSSA KO cells (Fig. 3E).

In order to determine whether the differential Pol II degradation across TC-NER deficient cells is specific for UV-induced damage, or also observed upon induction of other structurally different types of TBLs, we exposed cells to the mushroom-derived drug Illudin S, which induces TBLs that are recognized by TC-NER (46). Treatment with Illudin S increased Pol II degradation in UVSSA KO cells compared to CSA and CSB KO cells, similarly as was observed after UV-induced damage (Fig. 3F). Together this shows that lesion-stalled Pol II is not degraded in CSA and CSB KO cells, resulting in prolonged stalling of elongating Pol II on TBLs, while lesion-stalled Pol II is efficiently extracted from the chromatin by VCP resulting in its degradation in UVSSA KO cells.

### Residence time of lesion-stalled Pol II correlates with patient-specific mutations

To directly link the observed differences in UVSSA KO cells versus the CSA and CSB KO cells to the UV^S^S and CS disorders, we studied Pol II chromatin binding in cells expressing well-defined patient-specific CSA mutations that either give rise to CS or UV^S^S. We selected the W361C mutation causing UV^S^S (30) and two CS-causing CSA point mutations. W194C giving rise to CS-I, the classical severe form of CS, and A160T results in CS-III, a late onset form of CS with milder clinical features (25, 47). Based on the CSA-DDB1 structure, the CS-causing W194C and A160T mutations are predicted to interfere with the CSA fold, which is more severely disrupted by the W194C mutation. Both mutations interfere with the binding of CSA to DDB1 resulting in the inactivation of the CRL4^CSA^ ligase activity (38). In line, A160T prevents the incorporation of CSA into the CRL4^CSA^ complex (48). In contrast, the UV^S^S-causing W361C mutation severely impairs UVSSA recruitment, while leaving the CRL4^CSA^ activity intact (30). FLAG-tagged WT CSA, or FLAG-CSA harboring the patient-specific point mutations was expressed in MRC5^GFP-RPB1^ CSA KO cells. Full length expression of the recombinant constructs was determined by western blot (Fig. 4A). The TC-NER status of the cell lines was confirmed by testing UV sensitivity by colony formation. WT CSA expression rescued the UV-sensitivity in CSA KO cells. However, CSA mutants did not, consistent with a full TC-NER deficiency prevalent in CS and UV^S^S patients (Fig. 4B).

**Figure 4.**
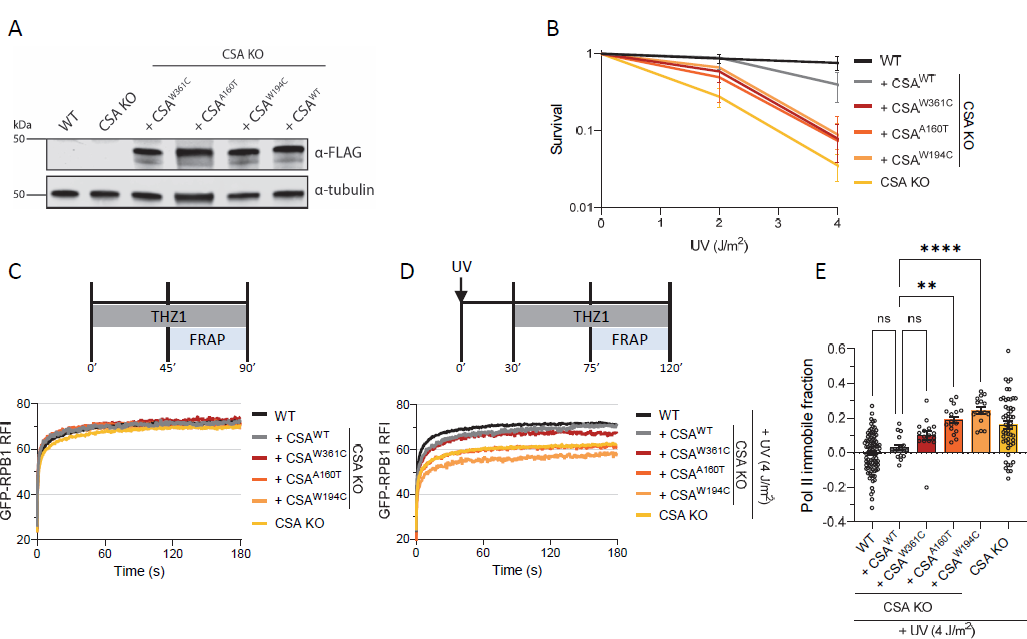
Pol II mobility correlates with patient-specific UV^S^S and CS mutations. A. Western blot of MRC5^GFP-RPB1^ WT and CSA KO cells complemented with WT FLAG-tagged CSA or FLAG-CSA harboring point mutations W361C, A160T or W194C. Exogenous expressed CSA is stained with FLAG antibody and tubulin is used as loading control. B. Relative clonogenic survival of the indicated cell lines in response to the specified UV doses. ± SEM, n ≥ 3. C. FRAP of GFP-RPB1 upon CDK7 inhibition with THZ1. GFP-RPB1 mobility of unperturbed WT and CSA KO cells is plotted in black and yellow, respectively, for comparison. Average RFI of n ≥ 16 cells per condition from at least 2 independent experiments. D. FRAP of GFP-RPB1 similar to (**C**), but after UV irradiation (4 J/m^2^) 30 min prior to CDK7 inhibition with THZ1. GFP-RPB1 mobility of UV-irradiated and THZ1 treated WT and CSA KO cells is plotted in black and yellow, respectively, for comparison. Average RFI of n ≥ 16 cells per condition from at least 2 independent experiments. E. Quantification of GFP-RPB1 immobile fraction from (**C**) and (**D**). n ≥ 16 cells per condition from at least 2 independent experiments. Ns = not significant, **P<0.01, ****P<0.001, ± SEM.

Next, we performed Pol II FRAP to assess Pol II chromatin binding in the cells expressing these patient-specific CSA mutants. In undamaged conditions no differences in Pol II chromatin binding were observed in the different CSA mutant expressing cell lines (Fig. 4C). Upon DNA damage induction followed by THZ1 treatment, Pol II chromatin binding in CSA^WT^ expressing cells was similar to WT cells, indicative of a full rescue of the prolonged Pol II stalling in CSA KO cells. In line with our hypothesis, CSA^W361C^ expressing cells showed a very similar Pol II mobility as CSA^WT^ expressing cells, while Pol II was markedly immobilized in CSA^A160T^ and CSA^W194C^ expressing cells, to almost a similar level as in CSA KO cells (Fig. 4D-E). By using mutations in the same protein either causing CS or UV^S^S we thus confirm that in UV^S^S cells, lesion-stalled Pol II is resolved, while the prolonged stalling of Pol II at TBLs is a specific feature of CS cells, and therefore contributes to the severe CS phenotypes.

## Discussion

This study shows that Pol II is differentially extracted from chromatin upon DNA damage in CS and UV^S^S cells and that impaired extraction of Pol II from TBLs is associated with increased severity of TC-NER related syndromes. We show that live cell imaging mobility studies of Pol II following inhibition of transcription initiation is a powerful setup to study the residence time of lesion-stalled Pol II. Using this approach, we observed a prolonged binding of Pol II to UV-damaged chromatin in CS cells, while Pol II was released from the chromatin in UVSSA and XPA KO cells to a similar degree as in TC-NER proficient cells. At the relatively low UV dose used, a fraction of 10-20% of all Pol II molecules is chromatin-bound at DNA damage in CS cells, representing 5.000-10.000 Pol II complexes that will form long-lasting roadblocks for transcription, thereby inactivating many genes. Additionally, these roadblocks will most likely impede other DNA transacting processes, or will result in transcription-replication conflicts, an important source of genome-instability (49). Such prolonged binding of Pol II at TBLs in CSB-deficient cells was confirmed in a recent study (50) showing retention of Pol II on CPDs while these complexes were resolved within 120 min after UV exposure in TC-NER proficient cells, consistent with our data also showing fully mobile Pol II 75-120 min after UV exposure in WT cells. In addition to Pol II, CSB was chromatin-bound for a prolonged time in CSA KO cells, indicating that in the absence of CSA, CSB is effectively recruited to lesion-stalled Pol II where it most likely remains as long as the lesion-stalled Pol II is present. In UVSSA KO cells, we could not study the CSB mobility by FRAP as in the absence of the deubiquitylating enzyme USP7 that is recruited by UVSSA, CSB is quickly degraded (data not shown), indicating that the CRL4^CSA^ is active at lesion-stalled Pol II in the absence of UVSSA.

We found that the degradation of lesion-stalled Pol II is almost completely absent in CS cells, in line with the lost CRL4^CSA^ activity in these CS cells. In contrast, in UVSSA KO cells ∼10% of lesion-stalled Pol II was degraded, in agreement with the ∼10-20% immobilization of lesion-stalled Pol II observed in CS cells, suggesting that almost all lesion-stalled Pol II is released from chromatin by degradation in UVSSA KO cells. This degradation in UVSSA KO cells is mainly caused by CRL4^CSA^ activity, while only a negligible contribution to Pol II degradation was observed for the E3 ligases NEDD4 and EloA, suggested to coordinate Pol II degradation as a last resort mechanism (45, 51, 52). In XPA KO cells, we observed, as expected, a Pol II degradation comparable to that of TC-NER proficient cells, consistent with the idea that Pol II has already been backtracked or released from the chromatin at the time of XPA recruitment.

We propose a model that explains the phenotypical difference between CS and UV^S^S (Fig. 5) in which Pol II stalling at TBLs happens in a similar manner irrespective of the TC-NER status but in which its processing happens differently. In WT cells the majority Pol II is released from the TBL through repair by TC-NER, without being directly degraded. Whether Pol II is released from the chromatin during the repair reaction, or is back-tracked and subsequently released upon transcription termination at the end of the gene upon repair, remains an important open question in the TC-NER field. In the absence of UVSSA, Pol II also does not remain bound at the lesion, even though there is no TC-NER, because Pol II can still be ubiquitylated by the CRL4^CSA^ E3 ligase. Ubiquitylated Pol II will be recognized by VCP, extracted from chromatin and subsequently degraded by the 26S proteasome. As a consequence, the lesion is made accessible to alternative repair factors. A likely candidate pathway to perform this task is GG-NER which efficiently recognizes and removes UV-induced 6-4-PPs. However, GG-NER is inefficient at recognizing CPDs (53), explaining the cutaneous photosensitivity of UV^S^S patients. When CSB or CSA are defective, the CRL4^CSA^ E3 ligase complex fails to ubiquitylate lesion-stalled Pol II, leading to a prolonged association of Pol II to the lesion and a complete shielding of UV-induced TBLs from other repair processes (6). In time, due to accumulating DNA damage, this will result in a gradual increase of lesion-stalled Pol II complexes. In XPA-deficient cells on the other hand, while the lesion is most likely accessible due to Pol II release, both TC-NER and GG-NER are impaired, thus eliminating the main pathways for removal of bulky DNA lesions. Only lesions that are not substrates for NER can still be repaired by other pathways, such as Base Excision Repair (BER) that repairs oxidative damage. This might explain the phenotype of the XPA-associated De Sanctis Cacchione (DSC) syndrome, which is milder than CS (54).

**Figure 5.**
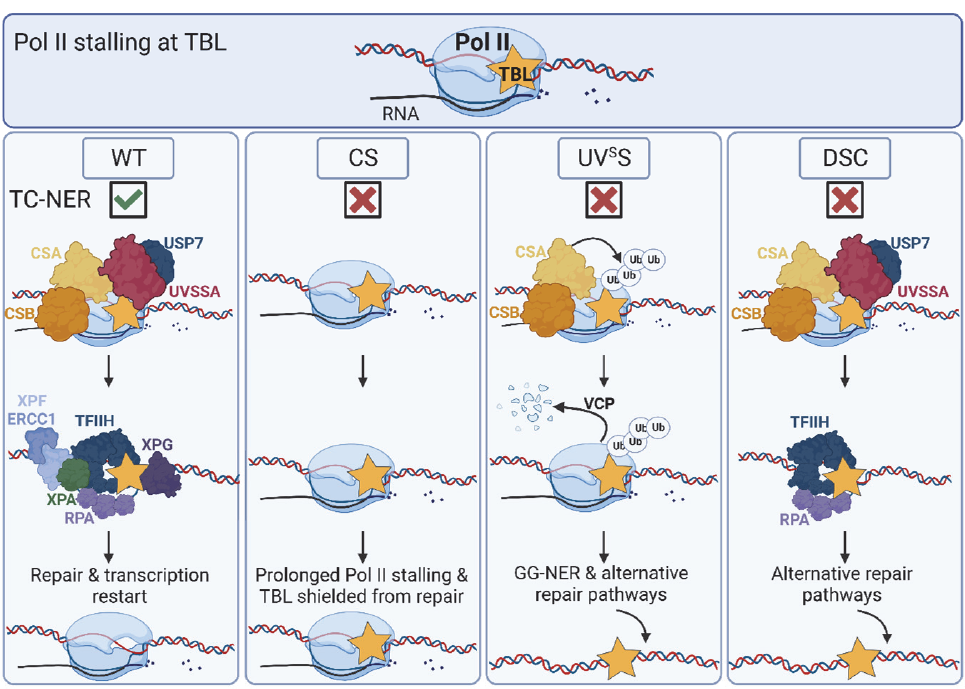
A. Model showing the fate of Pol II in WT, CS, UV^S^S, and XPA-associated De Sanctis Cacchione (DSC) cells. When elongating Pol II encounters a TBL it can have different outcomes. In WT and XPA KO cells Pol II will removed from the TBL during the repair reaction to allow repair of the lesion via TC-NER without being extensively degraded. However, in XPA KO cells TC-NER cannot be completed, but the lesion will be accessible for alternative repair pathways. In UV^S^S cells Pol II will be ubiquitylated in a CSA-dependent manner, removed from the lesion by the ubiquitin-specific protease VCP and subsequently degraded by the 26S proteasome, thereby exposing the lesion to other repair pathways that can eventually repair the damage. In CS cells Pol II will remain bound to the TBL creating a highly toxic persistent transcription block.

Our model also explains the striking differences in disease severity observed for different mutations in a single gene, as is the case for CSA. Mutations such as CSA^A160T^ and CSA^W194C^, affecting the E3 ligase function, will prevent the release of Pol II from chromatin and therefore cause a more severe phenotype. On the other hand, the CSA^W361C^ mutation, interfering with the recruitment of UVSSA but leaving the CRL4^CSA^ activity intact, will allow ubiquitylation and extraction of lesion-stalled Pol II and result in a milder disease phenotype. This model furthermore predicts that mutations in other TC-NER factors that impede Pol II ubiquitylation will result in CS-like phenotypes. For example, mutations in the recently identified TC-NER factor ELOF1 (19, 20), which drives Pol II ubiquitylation, are therefore expected to cause CS. In line with this, a mouse model in which the K1268 ubiquitylation site of Pol II is mutated features a CS-like phenotype, including growth retardation, progressive neurodegeneration and early death (16). Importantly, this model suggests that it is not the damage by itself which determines the severity of the TC-NER associated syndrome, but that it is the ability to remove Pol II from the lesion which determines the disease outcome. This indicates that unresolved TC-NER intermediates are more problematic for the cell than the residual TBL itself.

The apparent toxicity of unresolved TC-NER intermediates may explain the existence of multiple cellular orchestrated transcriptional responses to UV damage to prevent Pol II molecules from encountering TBLs. Recently, UV-induced degradation of promoter-bound Pol II was described (41), which prevents Pol II from going into productive elongation and thus stalling on lesions. Not only Pol II, but also TFIIH occupancy at promoters is reduced upon DNA damage, presumably to allow TFIIH bound at promoters to participate in TC-NER. This also leads to a reduced transcription initiation resulting in a global shutdown of transcription (55). Active transcription repression is also known to occur, as exemplified by the action of ATF3 (56, 57) or BMI1 and UBR5 (58) that transiently inhibit transcription on UV-damaged chromatin. These mechanisms to prevent further Pol II stalling are highly relevant in conditions where TC-NER is limiting, *i.e.* at high damage loads, to decrease the amount of persistently lesion-stalled Pol II molecules.

Transcription is an essential process in every cell. Therefore, it is interesting to note that CS patients are especially characterized by extreme neurological dystrophy. Due to the post-mitotic nature of neurons, other mechanisms that could result in clearing of lesion-stalled Pol II, such as DNA replication-mediated Pol II clearance, will not be active. Alternatively, neurons might be particularly sensitive to DNA damage-induced transcription stress because they depend on expression of relatively long neuronal genes (59). Accumulation of elongating Pol II on gene bodies in a gene-length dependent manner due to endogenous damage, has recently been found to cause transcription stress and contribute to normal aging (60). Therefore, long neuronal genes have a high likelihood of accumulating DNA damage that will block transcription, explaining why neurons are so sensitive to TC-NER deficiency.

Obviously, neurodegeneration associated with CS and DSC can hardly be explained by the prevalence of UV-induced CPDs and 6-4-PPs. The type of damage that causes neurodegeneration in NER disease is currently not exactly known, but aldehydes, alkylating agents or reactive oxygen species (ROS) likely contribute to the accumulation of DNA damage in neurons (3). For example, the high oxygen consumption and the higher levels of mitochondrial and metabolic activity in neurons (61) will result in increased ROS levels, that may saturate BER, thereby putatively contributing to DNA damage-induced transcription stress (61). Only TBLs that are inaccessible for alternative repair pathways, due to the size and placement of Poll II, will lead to Pol II accumulation on transcribed genes over time and will require CRL4^CSA^-dependent Pol II degradation. Other obstructions such as macromolecular complexes crosslinked to the chromatin will obviously not be completely shielded by the elongating Pol II and may therefore be removed without the need to release Pol II by degradation. However, this still needs to be experimentally tested. Alternatively, also in this situation the macromolecular elongation complex might need to be degraded to stimulate efficient repair of structurally different types of TBLs. This suggests that Pol II degradation is a crucial protective mechanism activated upon encounters with DNA damage to safeguard the essential transcription process. This is in line with the observation that in addition to the CRL4^CSA^ complex several other E3 ubiquitin ligases have been described to ubiquitylate Pol II, including NEDD4 (51), EloA (52, 62), BRCA1-BARD1 (63) and VHL (44, 64, 65). Future research should uncover under which exact conditions, e.g. specific types of TBLs (e.g. UV-induced damage vs. crosslinks) or specific tissue types (e.g. post-mitotic neurons vs. highly replicative cells), these might play an essential role to ubiquitylate Pol II to protect transcription fidelity.

## Methods

### Cell culture

Cells were cultured in DMEM (Gibco) and Ham’s F10 (Invitrogen) mixed 1:1 supplemented with 10% fetal calf serum (Biowest) and 1% Penicillin-Streptomycin at 37°C with 5% CO_2_. Generation of WT and TC-NER deficient GFP-RPB1 and CSB-mScarletI KI cell lines is detailed in **SI Materials and Methods**. For siRNA transfection, cells were transfected twice using Lipofectamine RNAi Max (Invitrogen) according to the manufacturer’s instructions 2 days prior to performing flow cytometry measurements. siRNAs were acquired from Horizon (non-targeting control: CAGACAAUCUUAUUACACA-3’, siElongin A 5’-GCACAGAACCCAGAGAAA-3’, siNEDD4 5’-AUGGAGUUGAUUAGAUUACAA-3’).

### Clonogenic survival assay

400-500 cells were seeded in 6-well plates in triplicate. The following day (MRC5) or 2 days later (HC116) cells were UV irradiated and allowed to grow for 7-10 days. Cells were fixed and stained with 50% methanol, 7% acetic acid, and 0.1% w/v Coomassie brilliant blue (Sigma). Colony numbers were determined using a GelCount colony scanner (Oxford Optronix). Average colony number for each UV dose was normalized to mock-treated conditions which were set to 1.

### Cell fractionation and Western blotting

In short, chromatin and supernatant fractions were separated in a series of centrifugation-based steps and DNA in the pellet fraction was degraded with benzonase. The procedure is described in detail in **SI Materials and Methods**. Proteins were separated on SDS Page gels, blotted overnight at 30V. Membranes were blocked in 5% skim-milk in PBS and stained with primary antibodies (see antibody list). Secondary antibodies were conjugated to IR-Dyes (Sigma) and detected using Odyssey CL or CLx infrared scanners (LiCor).

### Transcription Recovery assay

Transcription after indicated treatments was visualized by click-it based EU incorporation. The detailed procedure can be found in **SI Materials and Methods**.

### FRAP

FRAP was performed on a Leica SP5 confocal microscope or Leica SP8 using a HCX PL APO CS 63x, 1.40NA oil-immersion lens and LAS AF software. Fluorescence of GPF-RPB1 was detected using a 488 nm argon laser and CSB-mScarletI with a 561 nm laser. The detailed procedure can be found in **SI Materials and Methods**.

### Flow cytometry

Cells were harvested by trypsinization, washed in PBS and fixed in 1% Formaldehyde in PBS. Fluorescence levels were analyzed on a LSRFortessa Cell Analyzer (BD) equipped with FACSDiva Software (BD) and 10.000-20.000 cells were recorded per experiment. After exclusion of dead cells by granularity (SSC-A) and size (FSC-A), GFP levels were detected using a 488 nm laser and 530/30 filter. Flow cytometry data was analyzed using the FlowJo software (v.10.8.1) from BD Biosciences and used to determine the mean fluorescence intensity of GFP-RPB1. Fluorescence intensity was corrected and normalized to mock-treated fluorescence, which was set to 1.

## Acknowledgements

We thank the Optical Imaging Center for their support with microscopes and image analysis. This work is part of the Oncode Institute, which is partly financed by the Dutch Cancer Society. This study was supported by VICI Grant of Netherlands Organization for Scientific Research grant (VI.C.182.025).

---

**Supplementary Figure 1.**
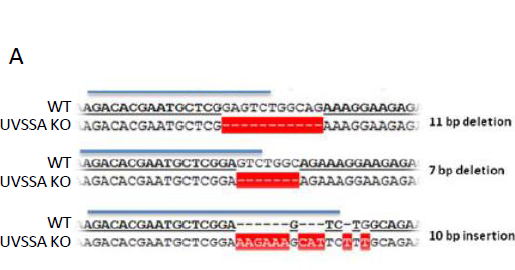
A. Genotyping results of MRC5^GFP-RPB1^ UVSSA KO cells used in this study. Position of the used gRNA is indicated by the blue line. Observed insertions, deletions and mismatches in the UVSSA KO cells are marked in red.

**Supplementary Figure 2.**
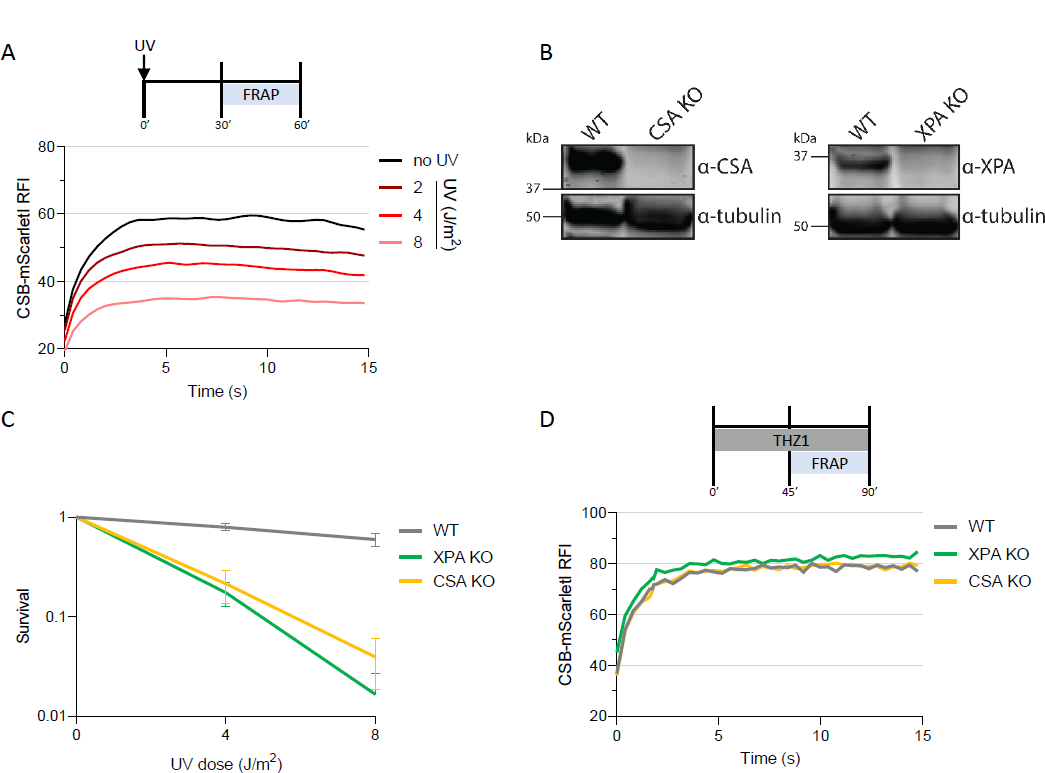
A. FRAP of CSB-mScarletI in HCT116^CSB-mScarletI^ showing UV-dose-dependent CSB immobilization. CSB-mScarletI was bleached in a strip across the nucleus and fluorescence intensity was measured every 0.4 sec for 16 sec 0-30 min after UV irradiation with the indicated UV doses from 3 independent experiments. B. Western blot analysis of the indicated HCT116^CSB-mScarletI^ WT and TC-NER KO cell lines using CSA and XPA antibodies. Tubulin was used as a loading control. C. Relative clonogenic survival of HCT116^CSB-mScarletI^ WT and TC-NER KO cells in response to the indicated UV doses. ± SEM, n ≥ 3. D. FRAP of CSB-mScarletI in HCT116^CSB-mScarletI^ WT and TC-NER KO cells as in (**A**), in unperturbed THZ1-treated cells. n ≥ 24 cells per condition from at least 2 independent experiments.

**Supplementary Figure 3.**
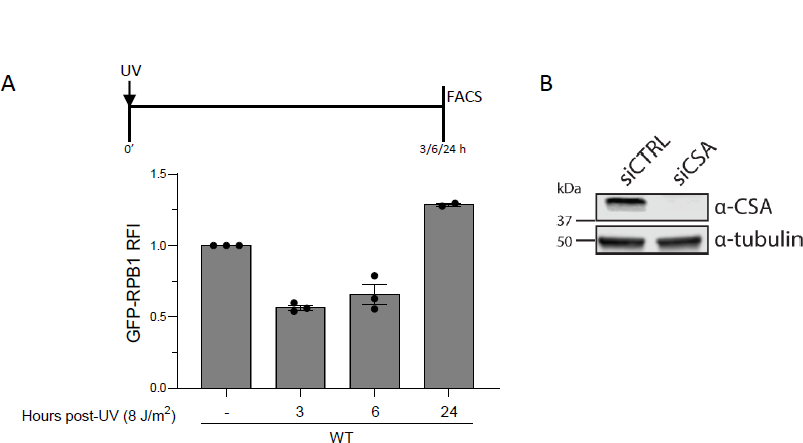
A. Pol II levels in MRC5^GFP-RPB1^ WT cells determined by flow cytometry 3 h, 6 h and 24 h after UV-irradiation (8 J/m^2^). GFP-RPB1 levels in UV-treated samples were normalized to mock-treated level which was set at 1. Columns represent average RFI of n ≥ 2 independent experiments ± SEM. B. Western blot analysis. MRC5^GFP-RPB1^ cells were stained with the indicated antibodies showing CSA-depletion with siCSA. SiCTRL was used as a non-targeting siRNA control. Tubulin was used as a loading control.

## SI Materials and Methods

### Cell culture and cell line generation

MRC5 (sv40) fibroblasts and HCT116 colorectal cells were cultured in DMEM (Gibco) and Ham’s F10 (Invitrogen) mixed 1:1 supplemented with 10% fetal calf serum (Biowest) and 1% Penicillin-Streptomycin at 37°C with 5% CO_2_.

HCT116 knockout (KO) cells were generated by transiently transfecting HCT116 osTIR1 CSB KI cells (19) with a pLentiCRISPR.v2 plasmid containing sgRNA targeting (CSA: (5’-GTCCGCACGCCAAACGGGTT-3’), or XPA: (5’-GTATCGAGCGGAAGCGGCAG-3’). Transfected cells were selected using 1 µg/ml Blasticidin (Invitrogen) for 7 days and single cells were seeded to allow expansion. Genotyping of single-cell clones was performed by immunoblotting (see antibody list).

MRC5 UVSSA KO clones were generated using a dual, doxycycline-inducible CRISPR/Cas9 vector system (iKRUNC, crRNA sequence: 5’-AGACACGAATGCTCGGAGTC-3’) and verified by TIDE analysis and sequencing of a subcloned PCR fragment of the genomic targeting locus using primers FW: (5’-CATTCTCCTGCCTCAATCTC-3’) and RV: (5’-CCTGTGCCTGGCATCTCTG-3’). MRC5 GFP-RPB1 knock-in (KI) TC-NER deficient cells were generated by targeting of the RPB1 locus as described in (41) in CSA, CSB, XPA and XPC KOs described in (42) and UVSSA KO.

pEGFP-N1 CSA^WT^ and CSA mutant plasmids were generated as previously described (48). The ORFs were amplified with Q5 polymerase (Invitrogen) using primers (5’-TCCTTCTTCATCACTGCTGC-3’) and (5’-TCACTTGTCGTCATCGTCTTTGTAGTCTCCTTCTTCATCACTGCTGC-3’) adding a FLAG-tag at the C-terminus and cloned into pEntr-D-Topo (Invitrogen) according to the manufacturer’s instructions. The CSA ORFs were transferred into a pLenti-CMV-puro vector (66) using Gateway ® LR cloning kit (Invitrogen) and transfected in HEK293T for lentivirus production. The lentivirus was subsequently transduced in MRC5^GFP-RPB1^ CSA KO cells, generating CSA^WT^, CSA^A160T^, CSA^W194C^ and CSA^W361C^ complemented cells which were selected with 5µg/ml puromycin.

DNA damage was introduced by UV irradiation or Illudin S treatment. For UV-induced damage cells were irradiated with UV-C light emitted by a 254 nm germicidal lamp (Philips) after washing with PBS. For Illudin S treatment, cells were incubated with 250 ng/mL (TOKU-E) for 24 h. The CDK7 inhibitor THZ1 (Xcessbio) was used at a concentration of 2µM and added to cells 30 min post-damage for 45-90 min. 100 µM cycloheximide (Sigma) was added to cells 2 h prior to UV irradiation or Illudin S treatment and kept for the duration of the whole experiment. Proteasome and VCP inhibitors were added to cells at the time of UV-irradiation and kept on the cells for the duration of the experiment. Proteasome inhibitor Mg132 (Sigma) was used at a concentration of 50 µM and VCP inhibitor NMS-873 (Selleck Chemicals) at 5 µM.

### Western blot

Samples were separated on Mini-PROTEAN TGX 4-15% precast gels (BIORAD) in 1X running buffer (1.44% w/v glycine, 0.3% w/v TRIS, 0.1% w/v SDS). Proteins were transferred onto an ethanol-activated PVDF membrane (0.45 µm, Merck Millipore) in 1X transfer buffer (0.3% w/v TRIS, 1.45% w/v glycine, 20% ethanol) at 30 V for 15 h at 4°C. Membranes were blocked in 5% skim-milk in PBS and stained with primary antibodies in PBS (see antibody list). Secondary antibodies were conjugated to IR-Dyes (Sigma) and detected using Odyssey CL or CLx infrared scanners (LiCor).

### Cell fractionation assay

Cells were seeded in 6 cm dishes, treated as desired and lysed for 30 min on ice in lysis buffer (30 mM HEPES pH 7.6, 1 mM MgCl2, 130 mM NaCl, 0.05% Triton X-100, 50 µM Mg132, cOmplete EDTA-free protease inhibitor (Roche) and Phosphatase inhibitor cocktail II (Sigma-Aldrich). Cell lysates were fractioned by centrifugation at 15.000 g for 10 min and the supernatant (cytoplasmic fraction) was collected in a separate Eppendorf tube. The cell pellet (chromatin-bound fraction) was first quickly washed with lysis buffer followed by a subsequent prolonged wash step in lysis buffer for 10 min on ice. After a second centrifugation step (15.000 g 10 min) the pellet was resuspended in 75 µL lysis buffer. Next, chromatin was degraded with 20 KU benzonase (Millipore) for 30 min on ice. 2x Laemmli sample buffer (Bio-Rad) was added to both pellet and supernatant fractions, boiled 5 min at 100°C and then loaded on an SDS gel.

### Transcription Recovery assay

Cells were grown in 6-well plates on glass coverslips and mock-treated or irradiated with UV (8 J/m^2^) 18 h or 2 h prior to addition of 200 µM 5’ethynyl uridine (EU, Jena Bioscience). Nascent RNA was labeled with EU for 1 h in Ham’s F10 (Invitrogen) + 1% PS + 10% dialyzed FCS supplemented with 20 mM HEPES buffer (Lonza). Cells were fixed in 3.6% Formaldehyde for 15 min and permeabilized in 0.1% Triton X-100 in PBS for 10 min. Cells were blocked in 1.5% BSA in PBS for 10 min and EU was fluorescently labelled via click-it chemistry for 1 h by incubation with 50 mM TRIS, 60 µM Atto-Azide-594nm (Atto-tech), 4 mM CuSO_4_*5H_2_O (Sigma) and 10 mM ascorbic acid (Sigma). After labelling cells were washed 3x in 0.1% Triton X-100 for 5 min, washed 2x in PBS and then DNA was stained with 100 ng/mL DAPI (Sigma) in PBS for 20 min to visualize nuclei. Finally, cells were washed 2x in PBS and mounted on glass slides using Aqua-Poly/Mount (Polysciences). Fluorescent images were acquired on a Zeiss LSM700 microscope using a 1.3 NA 40x oil-immersion lens and ZEN software. Image analysis was performed in ImageJ and DAPI signal was used to determine the nuclei of cells and thereby the region to measure EU signal. The integrated density of each UV-treated cell was normalized to the average integrated density of mock-treated cells, which was set at 1.

### FRAP

FRAP was performed on a Leica SP5 confocal microscope (for GFP-RPB1 expressing cells) or Leica SP8 (for CSB-mScarlet) using a HCX PL APO CS 63x, 1.40NA oil-immersion lens and LAS AF software. Fluorescence was detected using a 488 nm argon laser (GFP-RPB1) or 561 nm laser (CSB-mScarlet). A 512×32 pixel (GFP-RPB1) or 512×16 pixel-sized (CSB-mScarlet) strip was bleached across the nucleus of the cells using 100% laser power at 400 Hz for 1 (GFP-RPB1) or 2 frames (CSB-mScarlet). Fluorescence was measured at intervals of 0.4 sec for 25 frames pre-bleach and 450 frames post-bleach (GFP-RPB1) or 5 frames pre-bleach and 40 frames post-bleach (CSB-mScarlet). The fluorescence intensity of the nucleus in the bleached strip was background corrected to the pre-bleach fluorescence intensity outside of the nucleus within the same strip. Relative Fluorescence Intensity (RFI) was calculated by normalizing fluorescence values post-bleach to average values pre-bleach, which was set at 100. Immobile fractions (F_imm_) were calculated by comparing the UV-irradiated conditions to the mock-treated condition for each cell line using the following formula:

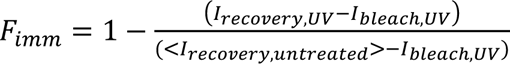

where I_recovery,UV_ is the average RFI of frames 250-450 post-bleach (GFP-RPB1) or frames 24-33 post-bleach (CSB-mScarlet) of each UV-treated cell, I_bleach,UV_ is the RFI directly after bleaching and < I_recovery,untreated_ > is the average RFI of frames 250-450 post-bleach (GFP-RPB1) or frames 24-33 post-bleach (CSB-mScarlet) of all cells in the untreated control cells.

### Antibody list

| Antibody | Host | Source | WB Dilution |
| --- | --- | --- | --- |
| CSA/ERCC8 | Rb | Abcam, ab240096 | 1:1000 |
| CSB/ERCC6 | Rb | Aboline, ABIN2855858 | 1:1000 |
| XPA | Rb | Genetex, GTX103168 | 1:1000 |
| XPC | Rb | Bethyl Laboratories, A301-122A | 1:2000 |
| Tubulin | m | Sigma Aldrich, B512 | 1:10000 |
| FLAG | Rb | Sigma Aldrich, F7425 | 1:1000 |
| RPB1-pSer2 | Rb | Abcam, ab5095 | 1:1000 |
| SSRP1 | m | Biolegend, 609702 | 1:5000 |
| Ku70 | Rb | Santa Cruz, sc-17789 | 1:1000 |

#### Secondary antibodies

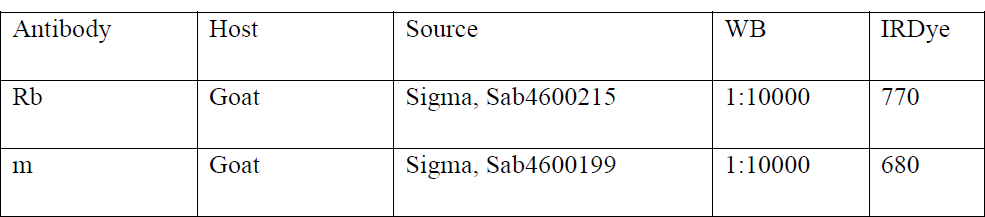

| Antibody | Host | Source | WB | IRDye |
| --- | --- | --- | --- | --- |
| Rb | Goat | Sigma, Sab4600215 | 1:10000 | 770 |
| m | Goat | Sigma, Sab4600199 | 1:10000 | 680 |

